# Early embryonic loss following intravaginal Zika virus challenge in rhesus macaques

**DOI:** 10.1101/2021.04.03.437254

**Authors:** Christina M. Newman, Alice F. Tarantal, Michele L. Martinez, Heather A. Simmons, Terry K. Morgan, Xiankun Zeng, Jenna R. Rosinski, Mason I. Bliss, Ellie K. Bohm, Dawn M. Dudley, Matthew T. Aliota, Thomas C. Friedrich, Christopher J. Miller, David H. O’Connor

## Abstract

Zika virus (ZIKV) is an arthropod-borne virus (arbovirus) and is primarily transmitted by *Aedes* species mosquitoes; however, ZIKV can also be sexually transmitted. During the initial epidemic and in places where ZIKV is now considered endemic, it is difficult to disentangle the risks and contributions of sexual versus vector-borne transmission to adverse pregnancy outcomes. To examine the potential impact of sexual transmission of ZIKV on pregnancy outcome, we challenged three rhesus macaques (*Macaca mulatta*) three times intravaginally with 1 × 10^7^ PFU of a low passage, African lineage ZIKV isolate (ZIKV-DAK) in the first trimester (∼30 days gestational age). Samples were collected from all animals initially on days 3 through 10 post challenge, followed by twice, and then once weekly sample collection; ultrasound examinations were performed every 3-4 days then weekly as pregnancies progressed. All three dams had ZIKV RNA detectable in plasma on day 3 post-ZIKV challenge. At approximately 45 days gestation (17-18 days post-challenge), two of the three dams were found to have nonviable embryos by ultrasound. Viral RNA was detected in recovered tissues and at the maternal-fetal interface (MFI) in both cases. The remaining viable pregnancy proceeded to near term (∼155 days gestational age) and ZIKV RNA was detected at the MFI but not in fetal tissues. These results suggest that sexual transmission of ZIKV may represent an underappreciated risk of pregnancy loss during early gestation.

## Introduction

Zika virus (ZIKV) emerged from relative obscurity five years ago to sweep through tropical and subtropical regions of the Western hemisphere. More than a million cases between 2015 and 2018 were reported in Pan American Health Organization (PAHO) regions alone (1). While ZIKV primarily causes mild febrile illness or asymptomatic infections in a majority of individuals, infection during pregnancy can result in a range of adverse outcomes including fetal loss and a constellation of birth defects now known as congenital Zika syndrome (CZS) (2–4). Human infection with ZIKV can occur following mosquito-borne, vertical, and sexual transmission (5–7). While mosquito-borne transmission from infected *Aedes* species mosquitoes is thought to be the most common route of infection in endemic areas, the contribution of sexual transmission in epidemics remains poorly understood, in part because during an outbreak, both transmission routes occur simultaneously and can be challenging to disentangle (8).

Sexual transmission of ZIKV was first documented in 2008 when a scientist working in Senegal became infected and, upon his return to the United States, infected his wife (9). Throughout the ZIKV outbreak in 2015 and 2016, additional sexually-transmitted infections were documented (10–14). The majority of sexually-transmitted cases in non-endemic areas are likely the result of infection of the primary cases during travel, followed by inadvertent transmission to the secondary cases upon returning home (7). As previously mentioned, sexually-transmitted ZIKV infections in endemic areas or areas experiencing active outbreaks are difficult to differentiate from mosquito-transmitted infections because there may be an individual risk of exposure by either route. Epidemiological data suggest that sexual transmission occurs primarily male-to-female through vaginal contact, even weeks after clinical symptom resolution, which suggests that sexual transmission of ZIKV does pose at least a theoretical risk to pregnant women (15). Furthermore, the ZIKV viral RNA (vRNA) load in human semen has been reported to range from the hundreds to tens of millions of copies per milliliter, with values as high as 3.98×10^8^ copies/ml reported (16– 18). The testes in particular, were found to be a ZIKV reservoir in animal models (19,20). In addition, studies have recently shown that intimate partners of household index cases are more likely to also be positive or show serologic evidence of ZIKV infection relative to other members of the same household (21).

Overall, we have limited information regarding the risk of ZIKV sexual transmission to pregnant women and their developing fetuses (14). Sexual transmission may be especially relevant during early pregnancy, since pregnancy can be inherently linked to unprotected sex. Likewise, studies have shown that other sexually transmitted ascending vaginal infections are associated with an increased risk of pre-term labor and other poor outcomes (22). Whether an ascending intravaginal ZIKV infection poses a higher risk to pregnancy than mosquito-borne infection is currently unknown. Pregnant women or women trying to become pregnant may be less likely to utilize condoms, a recommended strategy for the prevention of sexual transmission of ZIKV (23,24). Furthermore, a woman might not be aware of a pregnancy during early gestation and unfortunately, existing data suggest that the highest risk for developmental anomalies associated with ZIKV infection is during the first trimester, a critical developmental window (25–27). Additionally, ZIKV infection during pregnancy has also been associated with an increased risk for spontaneous abortion in both humans and nonhuman primates (28,29).

Animal models have played a critical role in improving our understanding of the natural history and pathogenesis of ZIKV. To-date, both murine and nonhuman primate (NHP) models have been utilized to examine aspects of sexual transmission of ZIKV (19,20,30,31). Studies in these models have shown persistent shedding of vRNA from the reproductive tract, infection of the female reproductive tract via a vaginal exposure route, and fetal effects as a result of vaginal exposure or sexual transmission in mice (20,30–39). Although studies in pregnant olive baboons have shown that intravaginal challenge with infected baboon semen during mid-gestation can result in productive maternal infection and vRNA detection in some maternal tissues and placentas, to date, studies in NHP have not shown clear evidence of vertical transmission associated with maternal ZIKV infection by the intravaginal route (33).

Because infection during the first trimester is associated with the highest risk for adverse pregnancy outcomes, and because unprotected sexual contact may be more likely during the first trimester, we designed a proof-of-concept study in which we challenged three gravid rhesus macaques (*Macaca mulatta*) intravaginally with ZIKV. Our goal was to investigate the potential impact of intravaginal ZIKV challenge during the first trimester on fetal and pregnancy outcomes and to develop a model for sexual transmission during early pregnancy.

## Methods

### Ethics Statement

All animal procedures conformed to the requirements of the Animal Welfare Act and protocols were approved prior to implementation by the Institutional Animal Care and Use Committee (IACUC) at the University of California, Davis. Activities related to animal care, housing, and diet were performed according to California National Primate Research Center (CNPRC) standard operating procedures (SOPs). SOPs for colony management and related procedures are reviewed and approved by the UC Davis IACUC.

### Study design

Female rhesus macaques (*Macaca mulatta*, N=3) were time-mated and identified as pregnant by ultrasound according to established methods (40). Prior to study assignment normal embryonic growth and development were confirmed by ultrasound. Females were challenged in the first trimester at approximately 30 days gestational age (trimesters divided by 55-day increments; term 165±10 days) with 1×10^7^ PFU ZIKV-DAK three times intravaginally at approximately two-hour intervals (**Table 1, Figure 1**). Pregnancies were monitored by ultrasound every 3-4 days post-challenge and then weekly from day 50 onward throughout the study period. Standardized parameters were assessed including fetal growth (greatest length then biparietal and occipitofrontal diameters, abdominal circumference, humerus and femur lengths) and structural development, amniotic fluid volumes and placental parameters, and compared to normal growth and developmental trajectories for the species (40). Dams were weighed at each sedation and blood samples were collected daily from day 3 through day 10 post-challenge, followed by bi-weekly until maternal plasma vRNA loads were undetectable, and then weekly until hysterotomy. Plasma and peripheral blood mononuclear cells (PBMCs) were isolated at all time points, and serum was collected on days 0, 14, and 24 post-challenge (dams 049-102 and 049-103), and on days 0, 14, 27, and 122 post-challenge for dam 049-101. Urine was collected by ultrasound-guided cystocentesis (∼1 ml) on days 7, 10, 14, 21, and 24 post-challenge (dams 049-102 and 049-103) and on days 7, 10, 14, 27, and 122 post-challenge for dam 049-101. Hysterotomies were performed for dam 049-102 and 049-103 at the end of the first trimester (post-detection of nonviable embryos by ultrasound) and near term (∼155 days gestational age) for dam 049-101.

**Table 1.**
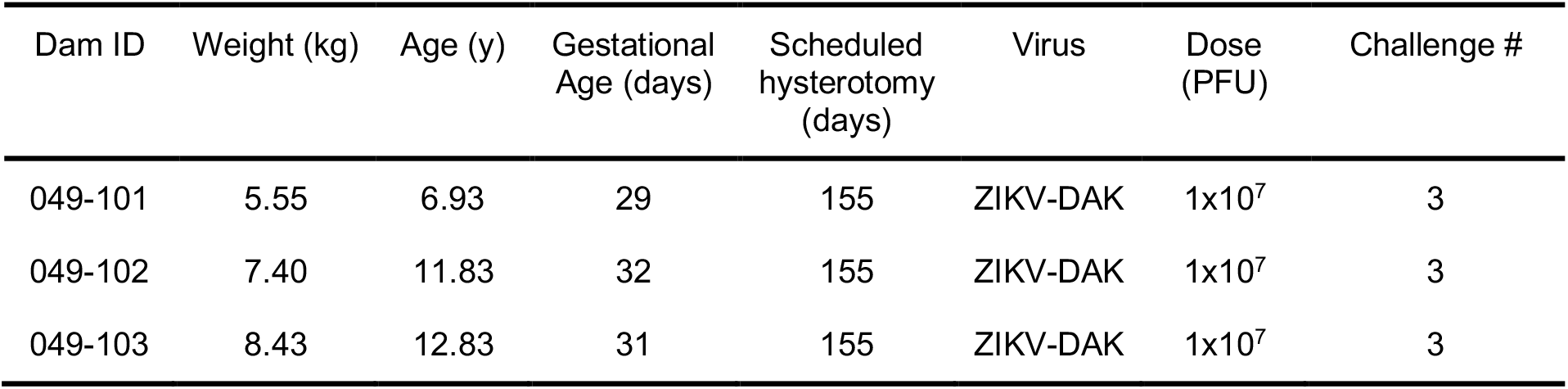
Dam information on day 0 of study.

**Figure 1.**
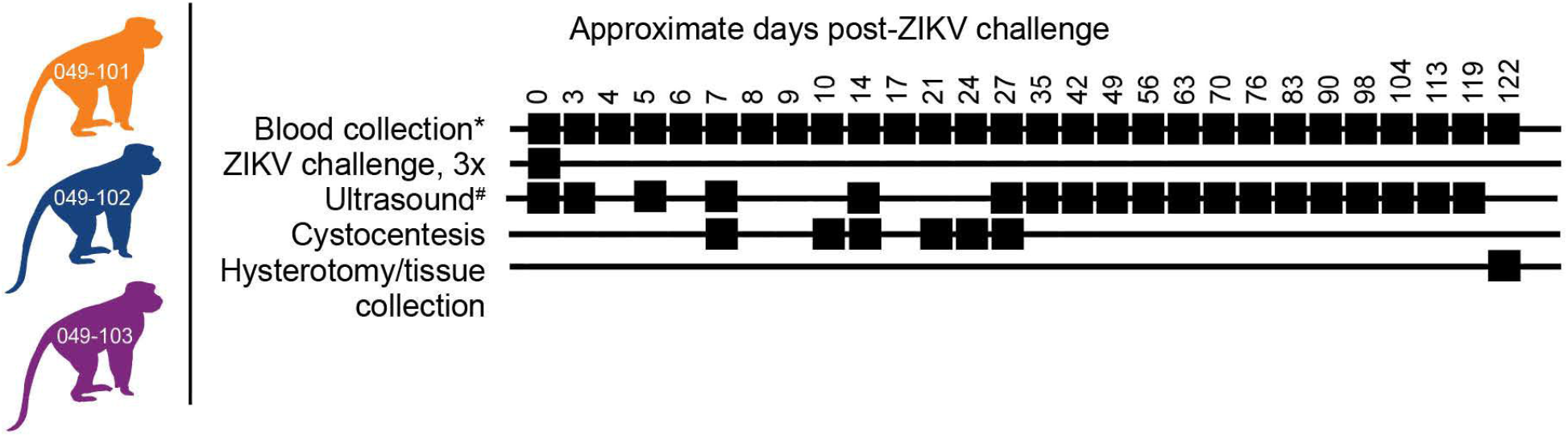
Study design. Three female rhesus macaques were time-mated, confirmed pregnant by ultrasound, and challenged intravaginally at ∼30 days gestational age with 1×10^7^ PFU ZIKV-DAK three times at two-hour intervals. Blood collection* denotes plasma and PBMC isolation at every sampling time point while serum collection was planned only on days 0, 14, 27, and 122 post-ZIKV challenge. Ultrasound^#^ denotes ultrasound imaging was performed every 3-4 days during early gestation, then weekly thereafter. Hysterotomies were originally planned for each animal at approximately 122 days post-ZIKV challenge.

### Virus challenge preparation and infection

ZIKV strain Zika virus/A.africanus-tc/Senegal/1984/DAK AR 41524 (ZIKV-DAK; GenBank: KX601166) was originally isolated from *Aedes luteocephalus* mosquitoes in Senegal in 1984. One round of amplification on *Aedes pseudocutellaris* cells, followed by amplification on C6/36 cells and two rounds of amplification on Vero cells, were used to prepare a master stock obtained from BEI Resources (Manassas, VA). Challenge stocks were prepared from this master stock by inoculation onto a confluent monolayer of C6/36 mosquito cells as described previously (41). Prior to administration, the ZIKV-DAK stock was diluted to 1×10^7^ PFU in 1 ml sterile saline and delivered via a 1 ml tuberculin syringe (37). Animals were inoculated three times intravaginally under ketamine sedation at approximately two-hour intervals using a previously described method (37).

### Blood processing and plasma vRNA loads

Plasma and PBMCs were isolated from blood placed in EDTA vacutainers and processed at 1500 RPM for 15 minutes according to standard protocols. Serum was isolated from whole blood collected into glass vacutainers without additives. Viral RNA was extracted from 300 µl plasma as previously described with a Maxwell 16 MDx instrument (Promega, Madison, WI) and evaluated using qRT-PCR (42). RNA concentration was determined by interpolation onto an internal standard curve of seven ten-fold serial dilutions of a synthetic ZIKV RNA segment based on Zika virus/Human/French Polynesia/10087PF/2013 (ZIKV-FP). The limit of quantification of the ZIKV qRT-PCR assay is estimated to be 100 copies vRNA/ml plasma or serum.

### Hysterotomy and tissue collection

Dams 049-102 and 049-103 were scheduled for hysterotomies in the late first trimester (nonviable embryos detected 3 days prior to hysterotomy). The hysterotomy for dam 049-101 was performed at approximately 155 days gestational age according to the original study design (**Figure 1**) and following established protocols (43). The gestational sac was removed for fetal tissue assessments, with a modified collection protocol for nonviable specimens (see below). For the fetus from dam 049-101 amniotic fluid, fetal blood, and fetal cerebrospinal fluid were collected, then fetal body weights and measures (biparietal and occipitofrontal diameters, abdominal and arm circumferences, hand and foot lengths, humerus and femur lengths, crown-rump length) were assessed. The left cerebral hemisphere and left eye were collected under aseptic conditions and shipped with cold packs to Wisconsin by overnight delivery for additional assessments (see below). Specimens collected for qRT-PCR for vRNA analysis included dura mater; right cerebral hemisphere (frontal, parietal, temporal, occipital lobes); cerebellum (right and left) and midbrain; right optic nerve; right eye (cornea, retina, sclera); spinal cord (cervical, thoracic, lumbar); right and left parotid glands, submandibular, and salivary glands; omentum; thymus; spleen; liver (right, left, quadrate, caudate lobes); pancreas; right and left adrenal glands and kidneys; right and left axillary and inguinal lymph nodes; diaphragm; tracheobronchial and mesenteric lymph nodes; right and left thyroids; trachea; esophagus; pericardium; aorta; right and left atria and ventricles; lung lobes (right and left; all lobes); reproductive tract including right and left gonads; urinary bladder; gastrointestinal tract (stomach, duodenum, jejunum, ileum, colon; meconium), skin, skeletal muscle, and bone marrow (**Table 2**). The placenta was weighed and assessed including disk measurements (primary and secondary for bidiscoid placentas; primary disk only for monodiscoid), umbilical cord and membrane insertion sites, blood vessel distribution, cut surfaces, and examined for the presence of infarcts. Decidua, membranes, umbilical cord, and multiple sections of the placental disks were collected. All specimens were quick frozen in triplicate over liquid nitrogen for qRT-PCR analysis or collected into RNAlater (cat# R0901, Sigma-Aldrich, St. Louis, MO). Multiple blocks of tissues were collected in histology cassettes fixed in 10% buffered formalin, embedded, sectioned (5-6 µm) and stained with hematoxylin and eosin (H&E) or used for *in situ* hybridization (ISH).

**Table 2.**
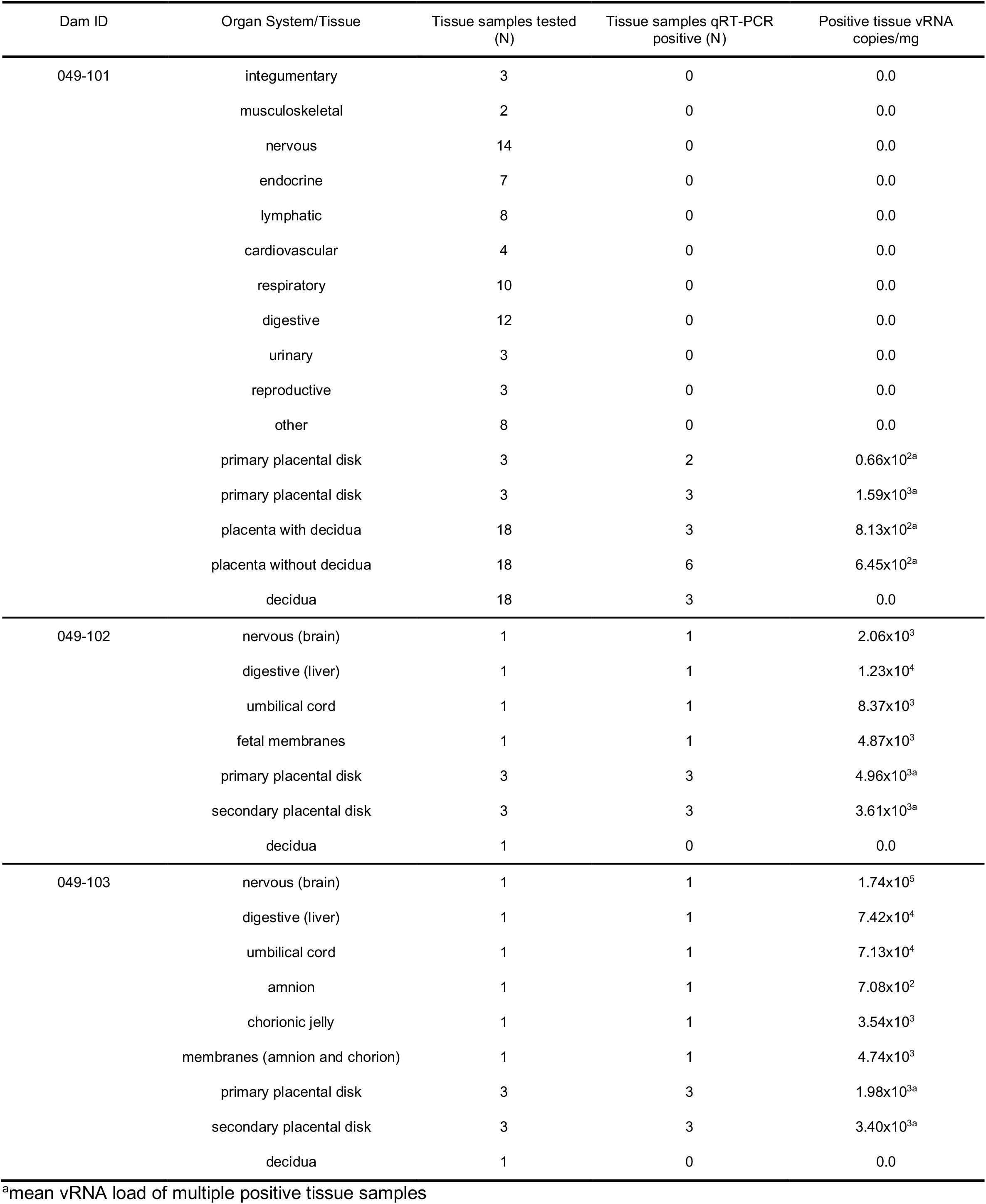
Fetal and maternal-fetal interface tissues collected at hysterotomy.

For dams 049-102 and 049-103 a modified collection was performed, consistent with the early developmental stage of the conceptus (**Table 2**). Decidua, membranes, umbilical cord, and multiple sections of the placental disks were collected as noted above.

Fresh samples collected from the 049-101 fetus (left cerebral hemisphere and left eye) were shipped with cold packs for additional assessments as noted above; the eye was analyzed by the Comparative Ocular Pathology Laboratory of Wisconsin (COPLOW). Placental tissues from all dams and tissues for the fetus from dam 049-101 were assessed as described previously in Koenig et al. (44).

### Tissue, urine, and amniotic fluid vRNA loads

Maternal-fetal interface (MFI) and fetal tissue vRNA loads were determined from approximately 20 mg of each tissue. ZIKV RNA was isolated from tissues using the Qiagen AllPrep DNA/RNA Mini Kit (cat# 80284, Qiagen, Germantown MD) using the QIAcube following the manufacturer’s protocol. Viral RNA was isolated from 140 µl maternal urine or amniotic fluid using the QIAmp Viral RNA minikit (cat# 52904, Qiagen, Germantown MD) following the manufacturer’s protocol. Following isolation, cDNA synthesis was performed using the Qiagen Sensiscript RT kit (cat# 205213, Qiagen, Germantown MD) according to the manufacturer’s protocol. Quantification of vRNA load was performed by real-time PCR using the Taqman amplification system and the QuantStudio 12 K Flex Real-Time PCR System (ThermoFisher Scientific, Grand Island, NY) as described previously (43). The estimated limit of quantification of the assay is 50- 100 ZIKV RNA copies/mg tissue (average = 75 copies/mg).

### *In situ* hybridization (ISH)

ISH probes against the ZIKV genome were commercially purchased (cat# 468361, Advanced Cell Diagnostics, Newark, CA). ISH was performed using the RNAscope® Red 2.5 kit (cat# 322350, Advanced Cell Diagnostics, Newark, CA) according to the manufacturer’s protocol. After deparaffinization with xylene, a series of ethanol washes, and peroxidase blocking, sections were heated with the antigen retrieval buffer and then digested by proteinase. Sections were then exposed to the ISH target probe and incubated at 40°C in a hybridization oven for two-hours. After rinsing, ISH signal was amplified using the provided pre- amplifier followed by the amplifier-containing labelled probe binding sites, and developed with a Fast Red chromogenic substrate for 10 minutes at room temperature. Sections were then stained with hematoxylin, air-dried, and mounted.

### Plaque reduction neutralization tests (PRNT)

Titers of ZIKV neutralizing antibodies were determined using plaque reduction neutralization tests (PRNT) on Vero cells (ATCC #CCL-81) with a cutoff value of 90% (PRNT_90_) (45). Neutralization curves were generated in GraphPad Prism (San Diego, CA) and the resulting data were analyzed by nonlinear regression to estimate the dilution of serum required to inhibit 90% Vero cell culture infection (45,46).

## Results

### Repeated intravaginal ZIKV challenge results in infection in pregnant macaques

All three dams had detectable ZIKV RNA in plasma by 3 days post intravaginal ZIKV challenge (**Figure 2**). ZIKV RNA loads peaked on day 5 for dams 049-101 and 049-102, and on day 6 for dam 049-103. Peak vRNA loads ranged from 1.57×10^4^ copies/ml for 049-101 to 1.30×10^5^ copies/ml for 049-103 (**Figure 2**). The latest detectable plasma vRNA load for animal 049-101 was on day 24 post-challenge (1.56×10^2^ copies/ml). Dam 049-103 had a detectable plasma vRNA load until day 14 (2.46×10^3^ copies/ml) but was negative on day 17 (the next time point samples were collected). Dam 049-102 was consistently positive for ZIKV vRNA until day 14, was negative on day 17, and then positive again on days 21 and 24 post challenge. Dam 049-102 was positive for ZIKV RNA in blood plasma collected at hysterotomy, the last time point sampled for the study. Overall, maternal plasma vRNA loads for dams 049-101, 049-102, and 049-103 were somewhat delayed compared to animals subcutaneously inoculated with French Polynesian or Puerto Rican ZIKV isolates in our previous studies, but were consistent in magnitude with previous observations (42,47,48). In addition, maternal plasma vRNA loads peaked within a time period similar to subcutaneously inoculated animals infected with the same ZIKV isolate (ZIKV-DAK) (49) (**Figure 2**).

**Figure 2.**
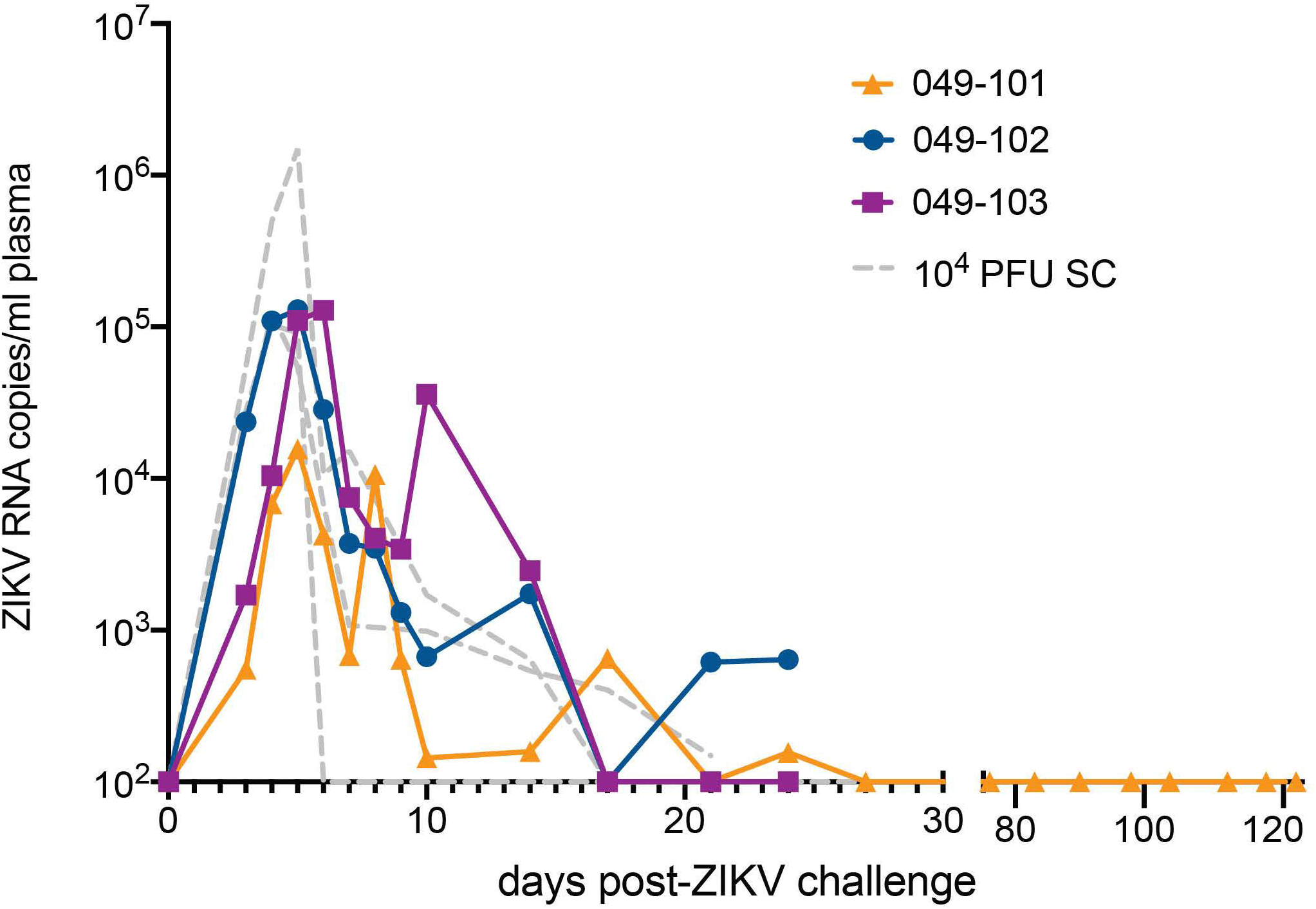
Intravaginal ZIKV challenge resulted in detection of vRNA in plasma for all three dams. The x- axis shows days post-ZIKV challenge. The y-axis starts at the estimated limit of quantification of the qRT- PCR assay (1×10^2^ copies/ml) and is shown as copies/ml plasma on the log scale. Plasma vRNA loads are displayed for dam 049-101 as orange triangles, for dam 049-102 as blue circles, and for dam 049-103 as magenta squares. For comparison, ZIKV plasma vRNA loads are also shown for three pregnant macaques subcutaneously (SC) inoculated with 1×10^4^ PFU ZIKV-DAK and are displayed as gray dashed lines and noted as 10^4^ PFU SC in the legend (49).

### Embryonic demise following intravaginal ZIKV infection during early pregnancy

Ultrasound examinations indicated that the embryos of dams 049-102 and 049-103 were nonviable at approximately 17-18 days post-challenge. Hysterotomies were subsequently scheduled and performed and each dam’s final blood and urine samples were collected (**Figure 3A**). Embryo and placental tissues from dams 049-102 and 049-103 were collected for vRNA analysis, histopathological assessment, and ISH. Dam 049-101’s pregnancy progressed normally and sampling continued until the study endpoint and near-term hysterotomy at approximately 155 days gestational age (**Figure 3A**). All fetal and placental measurements for 049-101 were recorded and were considered within normal limits for gestational age (**Table 3**) (40).

**Figure 3.**
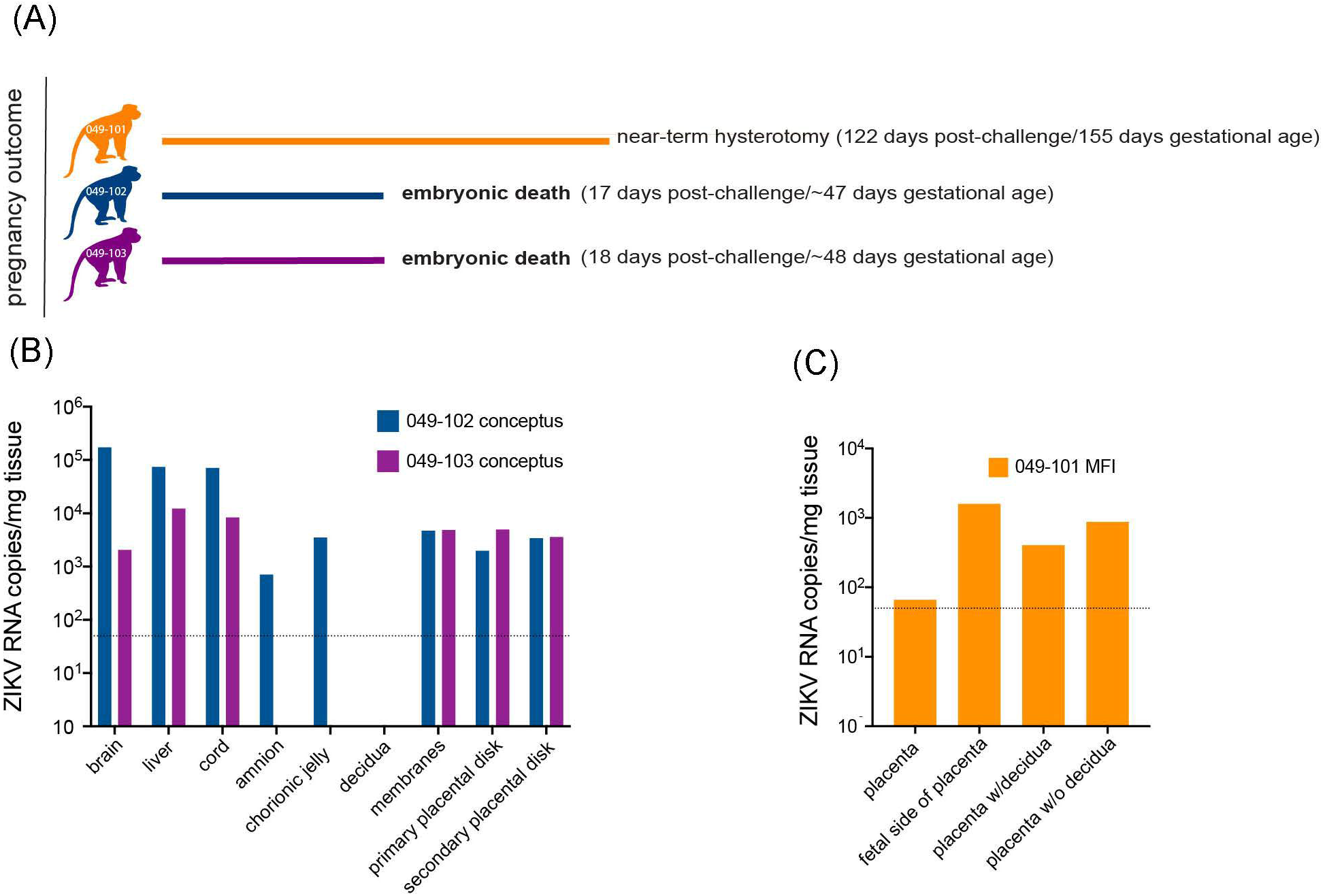
Pregnancy outcomes and maternal-fetal interface (MFI) and fetal tissue vRNA loads. **(A)** Pregnancy outcomes for three dams intravaginally inoculated 3x with ZIKV at approximately 30 days gestation. Two dams (049-102 and 049-103) were determined by ultrasound to have non-viable embryos at approximately 17-18 days post-ZIKV challenge. Hysterotomies and embryo and MFI tissue collections were performed 3 days after detection. Dam 049-101’s pregnancy continued until scheduled hysterotomy and extensive tissue collection at approximately 155 days gestational age. **(B)** Average ZIKV vRNA loads for positive embryo and MFI tissues collected at hysterotomy from dams 049-102 (blue) and 049-103 (magenta) following embryonic death at approximately 17-18 days post-ZIKV challenge. The dashed line represents the average of the estimated limit of detection (50-100 copies/mg, average: 75 copies/mg tissue) for the qRT-PCR assay. **(C)** Average ZIKV vRNA loads for positive MFI tissues collected at hysterotomy from dam 049-101 (orange) at approximately 122 days post-ZIKV infection. Fetal tissues were negative for ZIKV RNA by qRT-PCR. The dashed line represents the average of the estimated limit of detection (50-100 copies/mg, average: 75 copies/mg tissue) for the qRT-PCR assay.

**Table 3.** 049-101 fetal and placental measurements (∼155 days gestation, 122 days post challenge). Measurements were considered to be within normal limits by ultrasound and gross assessment (40). R = right, L = left.

| Tissue | Measure (mm) | Weight (g) |
| --- | --- | --- |
| whole body | 180.0 | 405.75 |
| biparietal diameter | 52.3 | - |
| head circumference | 186.0 | - |
| brain | - | 54.42 |
| cerebrum | - | 50.2 |
| cerebellum with midbrain | - | 3.31 |
| cerebellum without midbrain | - | 2.47 |
| r./l. eye | 13.6/13.8 | 1.34/1.35 |
| r./l. thyroid | - | 0.07/0.07 |
| thymus | - | 1.42 |
| spleen | - | 0.58 |
| liver | - | 12.08 |
| r./l. adrenal | - | 0.12/0.17 |
| r./l. kidney | - | 1.00/0.99 |
| lung lobes | - | 8.76 |
| r./l. testis | - | 0.06/0.06 |
| placenta | 145.0 x 85.0 | 159.19 |
| r./l. femur | 52.8/53.0 | - |

### MFI, fetal tissues, and amniotic fluid are ZIKV RNA positive in early embryos

ZIKV RNA was detected in the amniotic fluid from the conceptus of both dams 049-102 and 049-103 at 3.87×10^3^ and 7.38×10^3^ copies/ml respectively at the time of hysterotomy (subsequent to embryonic death). In addition, ZIKV RNA was detected in the brain and liver of both non-viable embryos, as well as in MFI tissues including the primary and secondary placental disks and membranes (amnion and chorion) (**Figure 3B**). The highest tissue vRNA burden was detected in the brain of the embryo from dam 049-102 (1.74×10^5^ copies/mg). ZIKV RNA was not detected in amniotic fluid collected from the fetus of dam 049-101 at hysterotomy. Although a large number of fetal and MFI tissues were assessed following hysterotomy, the presence of ZIKV RNA was only detected in a subset of sections of MFI tissues from 049-101 (**Table 2, Figure 3C**). The decidua from all three dams were negative for ZIKV RNA by qRT-PCR. Similarly, ZIKV RNA was not detected in the urine for any of the dams at any of the time points sampled. Overall, these results highlight the focal nature of ZIKV RNA detection in fetal and MFI tissues following infection during pregnancy. For a number of tissues, multiple samples were collected for vRNA analysis but ZIKV was only detected in a subset of those samples (**Table 2**).

### Pathological changes in placental tissues following intravaginal ZIKV infection are non-specific

Histopathological assessments of the placentas of dams 049-102 and 049-103 following embryonic demise showed generalized, non-specific mild necrosis (**Table 4**). In particular, the secondary placental disk from dam 049-102 showed significant necrosis for gestational age (tissues removed ∼50 days gestation) which was estimated to be approximately a week after embryonic death occurred. Placentas from both dam 049- 102 and dam 049-103 had minimal to mild multifocal villous mineralization. The primary placental disk of dam 049-102 showed moderate to marked intervillous hemorrhage and parenchymal ischemia. The placenta of dam 049-103 showed acute neutrophilic intervillositis and mild focal ischemia. In addition, the decidua from dam 049-102 showed some evidence of early decidual vasculitis. Similar to the placentas from the other two dams, the placenta of dam 049-101 showed mild, multifocal villous mineralization, findings which have previously been observed in control placentas. In addition, decidual tissue from dam 049-101 showed mild, multifocal muscularization of the decidual arteries. Overall, changes in the placental tissues were mild and not associated with any specific pathological processes. Assessment of fetal tissues from dam 049-101 showed normal brain and eye morphology with no identified lesions.

**Table 4.** Histopathological assessment of placental tissues from all animals and fetal tissues from 049- 101.

| Dam ID | Tissue | Findings |
| --- | --- | --- |
| 049-101 | primary placental disk | focally extensive hemorrhage within the basal plate; mild multifocal villous mineralization |
|  | decidua | mild multifocal persistent muscularization of decidual arteries |
|  | fetal brain | no pathological changes |
|  | fetal eye | no pathological changes |
|  | fetal lung | mild bilateral diffuse intra-alveolar squamous cells, similar to control |
| 049-102 | primary placental disk | moderate to marked intervillous hemorrhage and parenchymal ischemia with acute intervillous inflammation; minimal multifocal villous mineralization |
|  | secondary placental disk | significant necrosis for gestational age, nonviable embryo; not associated with specific pathologic process |
|  | decidua | early decidual vasculitis |
| 049-103 | primary placental disk | mild focal ischemia with coagulative necrosis and acute neutrophilic intervillitis; mild multifocal villous mineralization |
|  | secondary placental disk | changes in placental disks are mild and non-specific |
|  | decidua | minimal multifocal decidual necrosis with acute inflammation and two non-occlusive thrombi |

### ZIKV genomic RNA is detected in MFI tissues from demise cases

Tissue sections from decidua, primary placental disks, and secondary placental disks (bidiscoid placentas) were assessed by ZIKV ISH using RNAscope (see methods). ZIKV genomic RNA was detected in both the primary and secondary placental disks from dams 049-102 and 049-103 (**Figure 4**), but not from the primary placental disk from 049-101, nor any of the decidua sections from any of the pregnancies. The lack of ZIKV RNA in the decidua sections by ISH was consistent with the tissue vRNA assessment by qRT- PCR.

**Figure 4.**
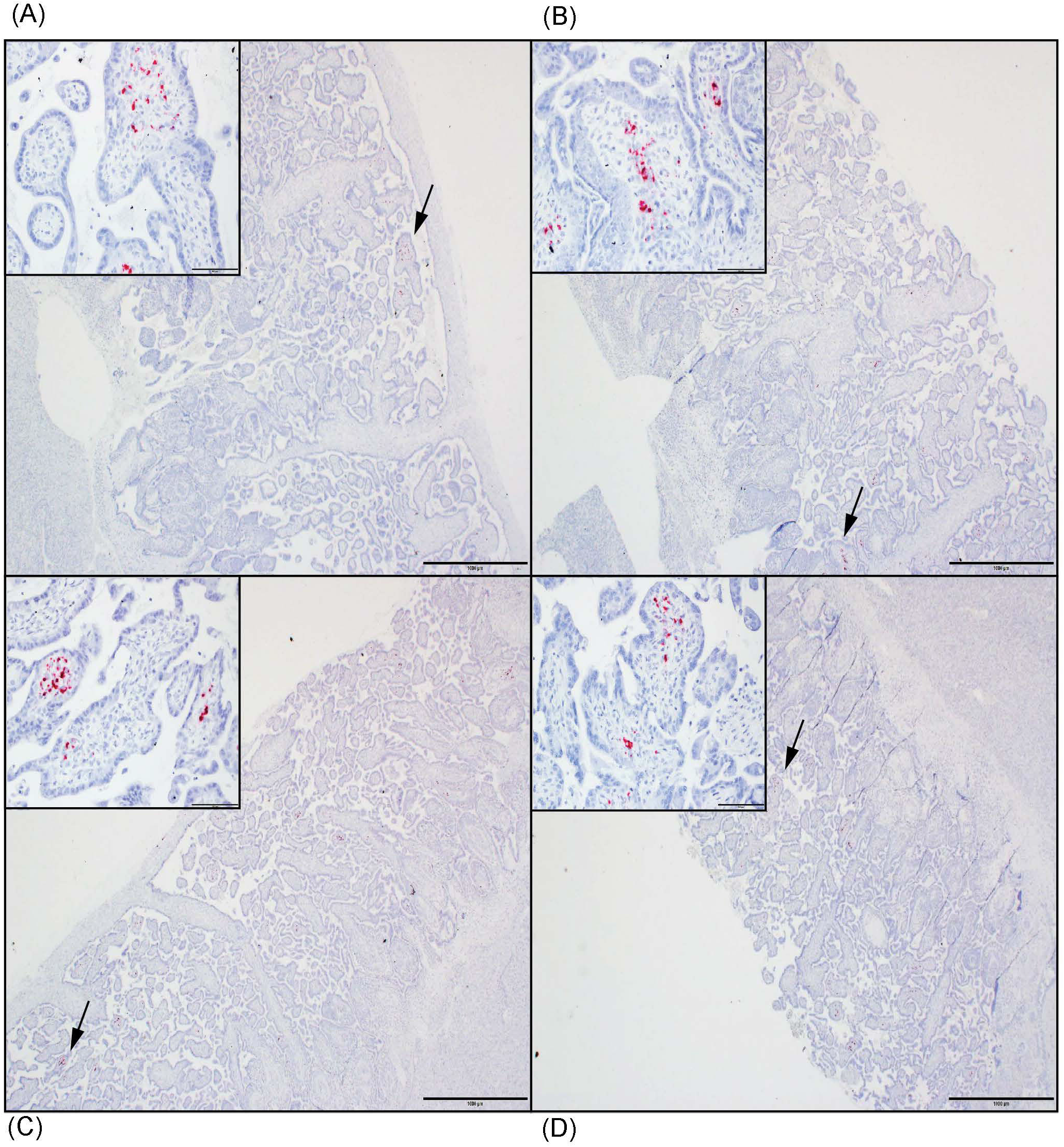
ZIKV genomic RNA was detected by *in situ* hybridization (ISH) in placental tissues collected from dams 049-102 and 049-103, but not from dam 049-101. For all images, red coloration is indicative of positive staining for ZIKV genomic RNA. Overall, positive staining is focal but visible in multiple areas. Insets show close-ups of the areas denoted by the black arrows in each larger panel. Representative images are shown of **(A)** primary placental disk from 049-103, **(B)** secondary placental disk from 049-103, **(C)** primary placental disk from 049-102, and **(D)** secondary placental disk from 049-102.

### Animals infected intravaginally with ZIKV during pregnancy develop neutralizing antibodies

Serum neutralizing antibody titers (nAbs) against ZIKV were evaluated for dams 049-102 and 049-103 on days 0, 14, and 24 post-challenge by 90% plaque reduction neutralization tests (PRNT_90_). Serum samples from 0, 14, 27, and 122 days post-challenge collected from dam 049-101 were similarly assessed. Samples collected on day 0 (pre-challenge) from all animals were negative for ZIKV nAbs. Neutralizing Ab titers above 1:10 are indicative of immunity against ZIKV. Serum collected on day 14 post challenge from all animals neutralized ZIKV-DAK at levels considered protective by PRNT_90_ (between 1:100 and 1:1000 for each animal). Serum collected on day 24 post-challenge from dams 049-102 and 049-103, and on day 27 post-challenge from dam 049-101 showed an increased neutralization response relative to baseline (day 0) and day 14 for each individual animal (**Figure 5**). By day 122 post-challenge, the ZIKV nAb response for animal 049-101 was lower than at days 14 or 27, but still demonstrated a strong protective response (PRNT_90_ titer approximately 1:300) (**Figure 5**). These results suggest that all animals developed a nAb response against ZIKV following intravaginal ZIKV challenge consistent with that previously noted for rhesus dams infected subcutaneously (42,47,48).

**Figure 5.**
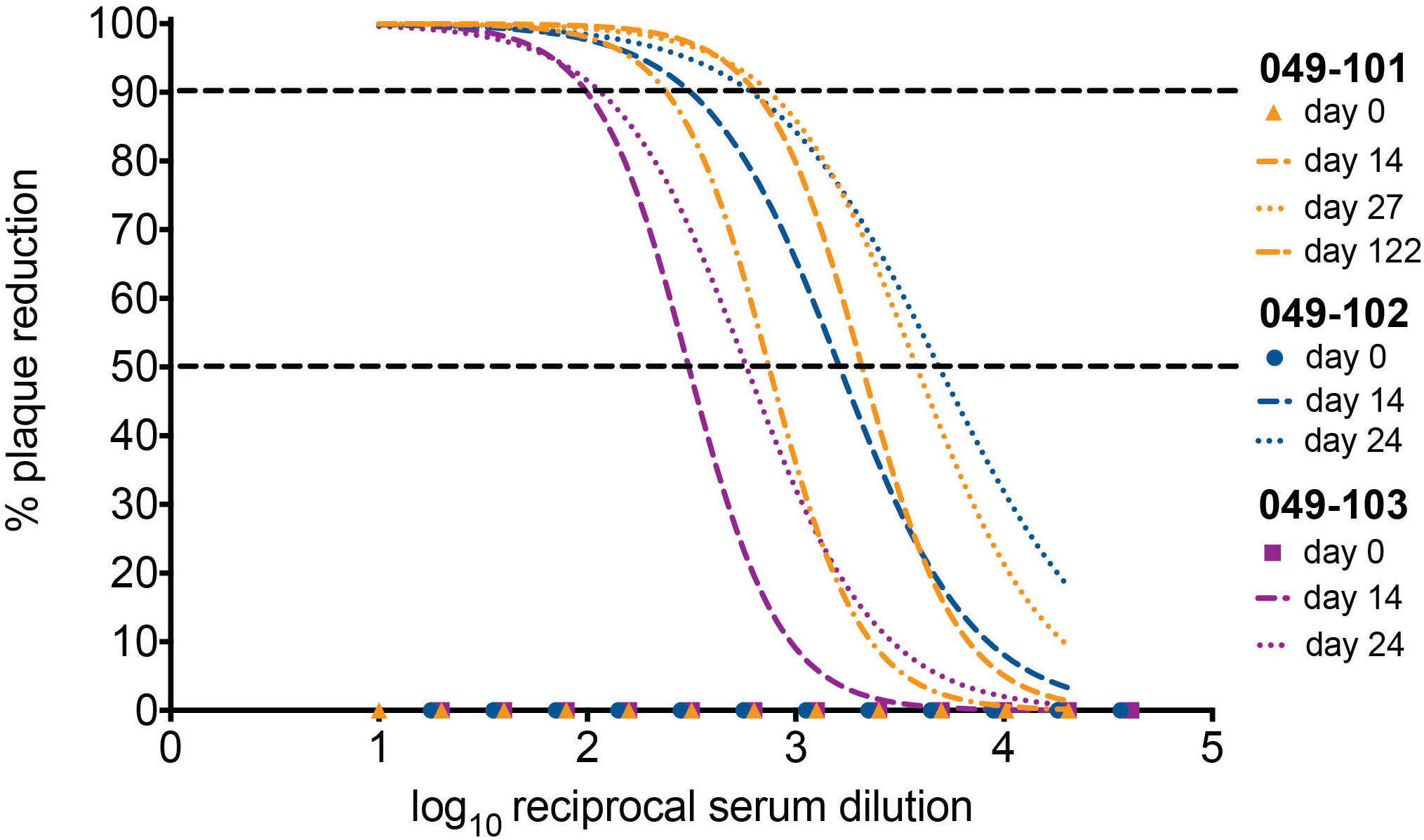
All three dams developed neutralizing antibodies (nAbs) against ZIKV as detected by PRNT_90_ following intravaginal ZIKV infection. The x-axis is the log10 reciprocal serum dilution and the y-axis is the percent plaque reduction for ZIKV-DAK. Day 0 for all animals is shown as symbols with 049-101 represented by orange triangles, 049-102 represented by blue circles, and 049-103 represented by magenta squares. Dashed gray horizontal lines indicate the PRNT_90_ and PRNT_50_ cut-offs respectively. Neutralization curves were generated using non-linear regression to estimate the dilution of serum required to inhibit 90% of Vero cell culture infection. Neutralization curves are shown for days 14 (dashed lines) and 24 (dotted lines) for dams 049-102 (blue) and 049-103 (magenta), and for days 14, 27, and 122 (dashed and dotted line) for dam 049-101 (orange).

## Discussion

Here we describe a proof-of-concept study that indicates intravaginal challenge with ZIKV during early pregnancy results in productive maternal infection and suggests that infection by this route can also result in embryonic demise. ZIKV RNA was detected at the MFI and in fetal tissues, as well as in the amniotic fluid from the pregnancies of dams 049-102 and 049-103, supporting a role for ZIKV in the adverse pregnancy outcomes for these animals. Although ZIKV was detected by qRT-PCR in the MFI tissues from dam 049-101, no vRNA was detected in fetal tissues directly. Interestingly, although vRNA was detectable in the placenta of dam 049-101 by qRT-PCR, it was not detected by ISH. Given the focal nature of ZIKV RNA detected in the placental tissue samples collected from dam 049-101, it is likely that the samples evaluated by ISH were simply from areas without vRNA present (**Table 2**). In order to assess transmission in these studies we intentionally avoided any intrauterine sampling to ensure no confounding variables. Because vRNA was not detected in any fetal tissues, our results may suggest that vertical transmission did not occur between dam 049-101 and the developing fetus. Alternatively, the results may suggest immunologic elimination of virus at later gestational ages as previously suggested by a study using direct fetal ZIKV inoculation (43). Our decision to challenge the animals in this study early in pregnancy (∼30 days gestation) was based on findings in humans suggesting that during the first trimester, ZIKV infection is associated with a higher risk of adverse fetal and pregnancy outcomes (27,50–53)(43)(27,50–53). In addition, we hypothesized that early pregnancy, possibly before a woman knows she is pregnant, may be a period of especially high risk for sexual transmission of ZIKV because precautions against this transmission route, such as condoms, may not be utilized (23,24). Overall, our results suggest that sexual transmission of ZIKV during early pregnancy may represent a significant risk for adverse outcomes.

Our results indicating early demise as a result of ZIKV infection are consistent with those described previously in a cross-center, cross-NHP species study (29). Interestingly, our finding that 2 of 3 (∼66%) pregnancies ended in nonviable embryos following intravaginal ZIKV infection during early pregnancy represents a higher rate of loss than the ∼26% previously reported for NHP (29). We acknowledge this loss rate is based on small animal numbers and could change as more animals are infected. Despite this higher rate compared to other NHP models reported to date, both near term and early gestation reflect periods of higher rates of spontaneous loss for macaques (54). While the loss rate reported in our study may be higher than the background rate of early loss in humans during the first trimester, data are very limited regarding the rate at which ZIKV-associated loss occurs in humans during the first trimester. A rate of around 11% was recently reported in a study during a period of epidemic transmission in Manaus, Brazil (55–58), although as noted, in many cases women may not be aware of an early pregnancy, thus the rate of loss could actually be higher. Additional studies with larger animal numbers will be necessary to determine the impact of the challenge dose, virus isolate, gestational age, and route of infection on pregnancy loss and how this relates to rates of spontaneous loss in early gestation.

Some limitations of this study include the use of a relatively high dose of ZIKV to inoculate the dams, the inclusion of multiple challenges over a short timeframe, and the small number of animals included in the study. The dose of inoculum chosen for this study is representative of the high end of the ZIKV vRNA range reportedly detected in human semen, which can be up to 100,000 times higher than that in blood (16–18). In part, this dose was also chosen due to the small number of animals included, our interest in the impact of intravaginal ZIKV exposure early in pregnancy, and the need to maximize chances of successful infection during early gestation. Previous studies in nonpregnant NHP have shown that intravaginal ZIKV inoculation results in successful infection after a single challenge approximately 33-75 percent of the time (31,37,59). In pregnant olive baboons, a single intravaginal inoculation mid-gestation with semen containing ZIKV (originating from French Polynesia or Puerto Rico) resulted in 4 of 6 animals developing detectable vRNA in blood, with an additional animal having detectable vRNA in blood after a second inoculation (33). This was the rationale for the choice to perform repeat challenges at two-hour intervals in this study: in order to maximize the likelihood of establishing a productive infection in our small cohort within a single day. We acknowledge that it is difficult to determine whether the inoculation route played a significant role in our observed outcomes or whether the cumulative inoculum dose, virus isolate, timing of infection, or some combination of these factors played a role in the observed outcomes. Future studies modeling sexual transmission should aim to determine which of these factors significantly impact pregnancy outcome.

We chose to utilize a low passage African ZIKV isolate (ZIKV-DAK) rather than a more contemporary isolate such as the commonly utilized PRVABC59 because, although it is also low passage, recent studies have suggested that this virus may have an attenuated phenotype and is not as pathogenic as ZIKV-DAK in mice (41,60). In addition, ZIKV was first isolated from a febrile rhesus macaque in the Zika Forest near Entebbe, Uganda in 1947 (61,62). Since that time, serologic and molecular (RNA or virus isolation) evidence of continued circulation in Africa has been intermittently reported in humans, animals, and mosquitoes (63–67). Prior to a report from Guinea-Bissau from 2016, during which an outbreak and subsequent identification of infant microcephaly cases was attributed to an African lineage virus, there were no reports of ZIKV impacting pregnancies and infant development in Africa (63,68). This has led to a number of hypotheses as to why, which includes, but is not limited to the following: widespread immunity in populations of childbearing age due to infection earlier in life; masking of ZIKV-associated adverse outcomes due to a high number of other, co-circulating pathogens in many populations, such as malaria; or embryonic loss during very early pregnancy simply unrecognized due to unknown status or inconsistent access to prenatal care (63,64,69). The data generated in this work supports the latter hypothesis of early loss. In reality, depending on the region, many of these factors could be playing an additive role in low and/or underreporting of ZIKV-associated pregnancy outcomes in Africa. Whether the early pregnancy losses observed in our study were due to increased pathogenicity of the African ZIKV isolate utilized relative to other isolates, the intravaginal route of infection, or both will require additional studies.

Many key questions remain with regard to understanding how different ZIKV geographic isolates may differentially impact pregnancy and fetal developmental outcomes. This study suggests that NHP models may be able to differentiate pregnancy outcomes between different isolates. Route of maternal infection may also play a role in pregnancy outcomes, at least in the case of NHP, as intravenous and intra-amniotic ZIKV infections during pregnancy have been associated with lower fetal survival rates across multiple studies compared to subcutaneous inoculation (29). As shown here, intravaginal infection may also lower survival rates in early pregnancy. Ultimately, our study was designed to balance all of the potentially influential factors previously mentioned within the constraints of a proof-of-concept study and the requirement for challenge and infection to occur during early pregnancy in order to evaluate this question.

Our results suggest that low passage, African lineage virus (ZIKV-DAK) has the potential to result in embryonic demise in rhesus macaques when infection occurs intravaginally and early in pregnancy. To our knowledge, this is the first NHP study to show clear evidence of vertical transmission of ZIKV following intravaginal infection, which has only previously been observed in mice (20,30,36). NHP, due to susceptibility without immune modulation, as well as having significant similarities to human pregnancy, may provide better approximations for human infections than other animal models (70). Furthermore, this is the first NHP study to show that African lineage ZIKV infection during pregnancy has the potential to result in severe fetal outcomes. Taken together, our results suggest that additional attention should be given to ongoing perinatal surveillance in African communities and to promoting awareness regarding the risks of sexual transmission of ZIKV in endemic areas.

## Author contributions

CMN, AFT, CJM, DHO designed the study.

AFT provided animal care, monitored the animals, and performed all sample collections.

AFT, CJM performed animal infections.

AFT, MLM, MIB, XZ, HAS, TKM, MTA, EKB analyzed samples.

CMN, DMD, JRR curated data.

CMN, JRR prepared the figures.

CMN prepared the initial manuscript draft.

All authors read, commented on, and edited the manuscript.

## Acknowledgements

We thank the research, animal care, and veterinary care staff at the California National Primate Research Center for caring for the animals and supporting this study amid the emerging SARS-CoV-2 pandemic. We also thank DHHS/PHS/NIH for providing the supplemental funding for R01 A1116382-01A1 that allowed this work to be completed. These studies were also supported by the California National Primate Research Center base operating grant #OD011107. *In vivo* imaging was performed with instrumentation funded by an NIH S10 grant #OD016261.

